# Neuroanatomical subtypes of Long COVID reveal distinct cognitive profiles alongside longitudinal brain changes

**DOI:** 10.64898/2026.07.30.741878

**Authors:** Angel Omar Romero-Molina, Betsabe McCord, Juan Fernandez-Ruiz, Adrian M. Owen, Carlos R. Hernandez-Castillo

**Affiliations:** Faculty of Medicine, National Autonomous University of Mexico, Mexico City, Mexico, 04360; Faculty of Computer Science, Dalhousie University, Halifax, Canada, NS B3H 1W5; Department of Physiology and Pharmacology, Western University, Ontario, Canada, ON N6A 5C1; Department of Psychology, Western University, Ontario, Canada, ON N6A 5C2

**Author notes:** Corresponding: Dr. Carlos R. Hernandez-Castillo Contact.

**Keywords:** Long COVID, magnetic resonance imaging, multilayer networks, Cluster analysis, Brain atrophy, neuroanatomical heterogeneity

## Abstract

**Background:** Long COVID is characterized by persistent symptoms following SARS-CoV-2 infection, including cognitive deficits among its most disabling manifestations. However, their neurobiological basis is poorly understood. Previous neuroimaging studies reported structural brain alterations, but few have integrated cognitive assessment, network-based analyses, and longitudinal imaging to identify neurobiological subtypes.

**Methods:** We studied 42 individuals with persistent post-COVID cognitive symptoms and 14 matched healthy controls using cognitive assessment and structural MRI. Multilayer brain networks were constructed from regional morphometric measures, and hierarchical clustering identified patient subgroups. Cross-sectional and longitudinal comparisons of brain structure, cognition, and function were performed, with exploratory brain–behavior correlations over six months.

**Results:** Clustering identified two subgroups with distinct structural patterns. Compared with controls, one subgroup showed reduced gray matter density in cerebellar lobules VIIIa/VIIIb and the putamen (p < 0.05, TFCE-corrected), whereas the other showed no significant alterations. Cognitive differences between clusters did not survive multiple-comparison correction; however, several measures showed medium-to-large effect sizes (d = 0.70–1.01), suggesting meaningful cognitive differences requiring confirmation in larger cohorts. Longitudinal analyses revealed increased medial frontal gray matter density (p < 0.05) associated with visuospatial/executive performance.

**Conclusions:** These exploratory findings suggest that post-COVID cognitive symptoms are associated with heterogeneous neurobiological profiles rather than a uniform pattern of impairment. Structural alterations involving cerebellar, striatal, and frontal regions may reflect distinct neuroanatomical phenotypes. Longitudinal findings suggest medial frontal structural reorganization with functional relevance. These findings support data-driven stratification for characterizing neurobiological heterogeneity in Long COVID and provide a foundation for future hypothesis-driven studies.

## 1. INTRODUCTION

The COVID-19 pandemic has left a significant proportion of individuals with persistent symptoms, collectively known as Long COVID. Among these, cognitive and neurological manifestations (commonly referred to as “brain fog”) are particularly disabling, affecting memory, attention, executive function, and processing speed [1]. A recent systematic review and meta-analysis documented that cognitive difficulties, especially impairments in memory, attention, and executive functions are highly prevalent in Long COVID and frequently co-occur with other neuropsychiatric symptoms, including anxiety and depression [2]. Importantly, evidence indicates that these abnormalities may emerge independently of the severity of the acute infection [3,4], suggesting that persistent cognitive deficits are not limited to individuals with severe initial disease. Collectively, these impairments substantially affect daily functioning and quality of life [5,6].

Neuroimaging studies have identified structural brain alterations in individuals with Long COVID, including reduced gray matter volume in frontal, temporal, occipital, and cerebellar regions, as well as disruptions in white matter microstructure [3,7,8]. Several of these alterations have been associated with objective cognitive deficits, supporting the existence of a neural substrate underlying persistent cognitive symptoms after SARS-CoV-2 infection [5,9]. However, the anatomical distribution and magnitude of these findings have varied considerably across studies, suggesting that Long COVID may not be characterized by a single pattern of brain involvement.

Despite growing evidence of cognitive and neurobiological abnormalities, a consistent clinical and neuroanatomical profile of Long COVID has yet to emerge. One possible explanation is that Long COVID represents a heterogeneous condition comprising biologically distinct subgroups rather than a single syndrome. Supporting this notion, symptom-based clustering studies have identified distinct clinical phenotypes characterized by different constellations of cognitive, neuropsychiatric, and peripheral nervous system symptoms [10]. More recently, imaging-based approaches have revealed divergent patterns of structural brain involvement, including subgroups characterized by predominantly cortical or cerebellar alterations despite similar cognitive outcomes at the group level [9]. Together, these findings suggest that heterogeneity may be a fundamental feature of Long COVID. Nevertheless, the neurobiological basis of this heterogeneity remains poorly understood.

Multilayer network analysis provides a promising framework for investigating this heterogeneity by integrating complementary sources of neuroimaging information within a unified representation of brain structure [11]. Unlike conventional single-modality approaches, multilayer models preserve the unique contribution of each imaging modality while simultaneously capturing interactions across modalities [12]. By characterizing large-scale patterns of brain organization rather than isolated regional alterations, this approach may facilitate the identification of biologically meaningful subgroups within heterogeneous clinical populations [13]. Consequently, multilayer network analysis represents a particularly suitable framework for exploring neuroanatomical variability in Long COVID and for identifying patient subtypes that may not be detectable using traditional neuroimaging analyses.

In the present study, we used multilayer network features derived from multimodal structural MRI to identify data-driven subgroups of Long COVID patients with persistent cognitive symptoms. We hypothesized that distinct neuroanatomical patient subgroups could be identified, reflecting the underlying heterogeneity of Long COVID. We then characterized these subtypes in terms of structural brain alterations, cognitive performance, health-related quality of life, and longitudinal trajectories over a six-month follow-up period. Finally, we investigated associations between structural brain changes and cognitive outcomes to assess the clinical relevance of the data-driven subtypes identified in this study. By moving beyond symptom-based classification, this approach aims to provide a more objective characterization of neuroanatomical heterogeneity in Long COVID.

## 2. MATERIALS AND METHODS

### 2.1 Participants

A total of 42 patients with post-COVID cognitive symptoms and 14 healthy controls matched for age, sex, and education level were included in this study (Figure 1). Inclusion criteria for the patient group were: (1) a confirmed history of SARS-CoV-2 infection verified by RT-PCR or antigen testing; (2) self-reported persistence of clinical and cognitive symptoms at the time of recruitment; and (3) absence of prior neurological or major psychiatric disorders. Exclusion criteria for all participants included contraindications to MRI scanning and a history of substance abuse.

**Figure 1.**
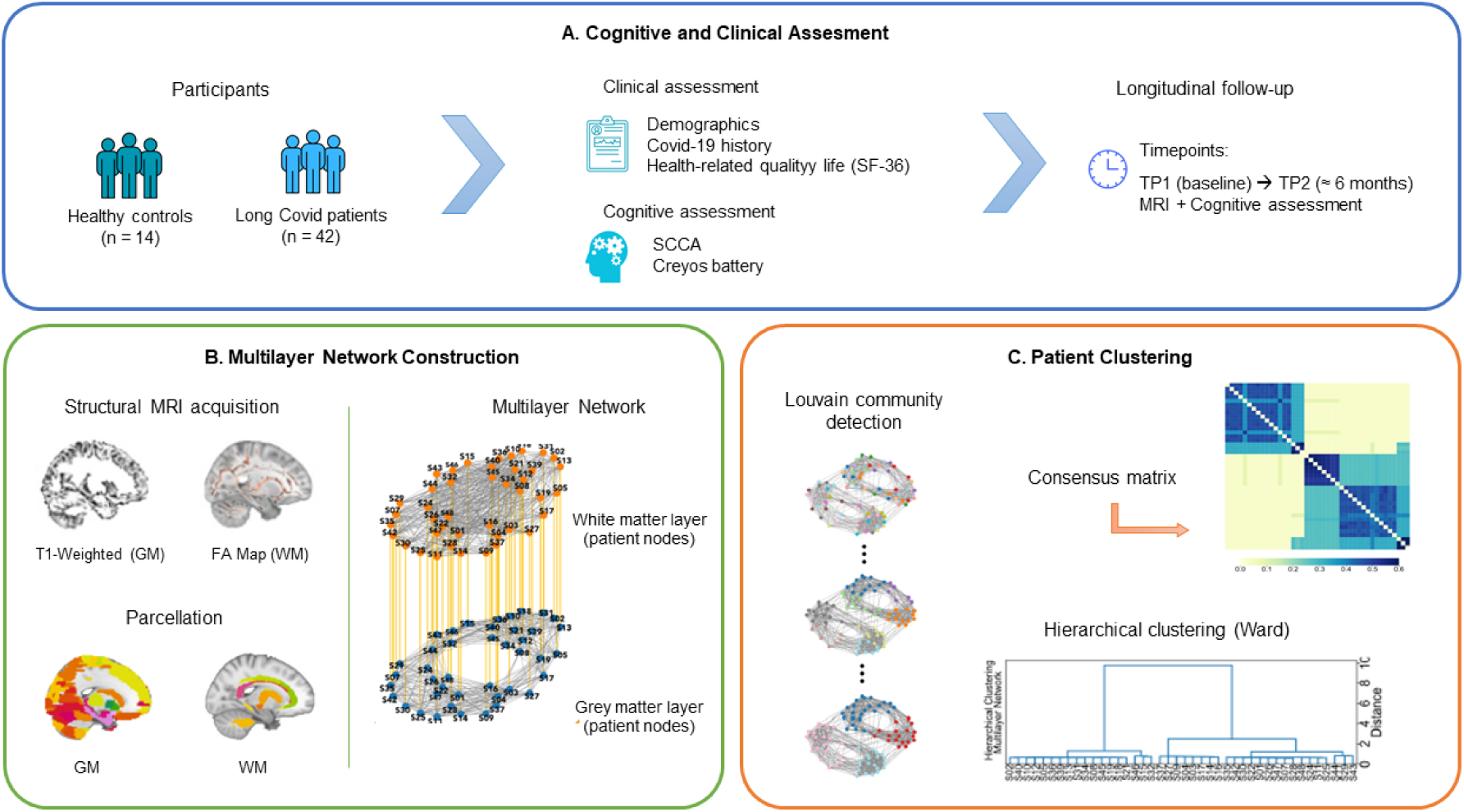
Overview of the study design and analytical pipeline. (A) Study design and participant assessment. Healthy controls and individuals with persistent Long COVID underwent structural MRI, comprehensive cognitive evaluation (SCCA and Creyos), clinical assessment, and health-related quality of life evaluation (SF-36). Longitudinal follow-up (TP2) was performed approximately six months after baseline (TP1). (B) Multilayer network construction. Structural MRI data wereprocessed to obtain gray matter (T1-weighted MRI) and white matter (fractional anisotropy) measures. After brain parcellation, tissue-specific structural information was integrated into a multilayer network in which each layer represented a structural modality and nodes corresponded to individual patients. (C) Clustering workflow. Patient subgroups were identified using Louvain community detection on the multilayer network. Consensus clustering was subsequently applied to generate a consensus matrix, followed by agglomerative hierarchical clustering (Ward’s method) to obtain the final patient subgroups used for subsequent structural, cognitive, and longitudinal analyses.

All participants underwent baseline cognitive, clinical, and neuroimaging assessments (Timepoint 1, TP1). In addition to cognitive testing, participants completed the 36-Item Short Form Health Survey (SF-36) to assess health-related quality of life across physical and mental domains. A follow-up assessment (Timepoint 2, TP2) was conducted approximately six months after baseline, during which 32 of the original 42 patients completed the longitudinal MRI assessment.

All participants provided written informed consent prior to participation. The study was approved by the research ethics boards of the Nova Scotia Health Authority and Dalhousie University and was conducted in accordance with the Declaration of Helsinki.

### 2.2 Cognitive assessment

All participants underwent a comprehensive cognitive assessment at baseline (TP1). Patients with Long COVID additionally completed the same cognitive battery at a six-month follow-up (TP2). Two complementary instruments were used to capture both clinically relevant cognitive dysfunction and performance-based cognitive abilities (Figure 1).

First, the Cerebellar Cognitive Affective Syndrome Scale (CCAS-S) was administered as a brief clinical screening tool designed to detect cognitive and affective disturbances associated with cerebellar dysfunction [14]. The CCAS-S assesses multiple cognitive domains, including executive functions, verbal fluency, visuospatial processing, and affective modulation, and yields both domain-specific scores and a global performance profile.

Second, participants completed the Creyos Cognitive Battery (formerly Cambridge Brain Sciences), a computerized, performance-based battery comprising 12 tasks that evaluate a broad range of cognitive domains, including working memory, attention, executive control, reasoning, and visuospatial abilities [15,16]. The tasks administered were Spatial Span, Digit Span, Paired Associates, Monkey Ladder, Grammatical Reasoning, Odd One Out, Double Trouble, Rotations, Feature Match, Token Search, Spatial Planning, and Interlocking Polygons.

Raw scores were extracted for all cognitive measures and used in subsequent analyses. Healthy controls completed the cognitive assessment at baseline only, whereas patients with Long COVID completed both baseline and six-month follow-up evaluations.

### 2.3 MRI acquisition and preprocessing

Structural MRI data were acquired for all participants at baseline (TP1) and for a subset of patients at six-month follow-up (TP2) using a 3 T MRI scanner (Discovery MR750, GE Healthcare, Waukesha, WI) equipped with a 32-channel head coil (MR Instruments, Inc., Minneapolis, MN, USA).

T1-weighted images were acquired for anatomical reference (voxel size: 1 × 1 × 1 mm; resolution: 184 × 224 × 224 voxels; echo time: 1.9 ms; repetition time: 4.4 ms; inversion time: 450 ms; flip angle: 9°). Diffusion-weighted images (DWI) were subsequently acquired using a single-shell protocol at b = 1000 s/mm² in 60 directions, with seven interleaved volumes at b = 0 s/mm² (voxel size: 2 × 2 × 2 mm; resolution: 108 × 108 × 77 voxels; echo time: 66 ms; repetition time: 8 s; flip angle: 90°), followed by eight reverse-phase images acquired at b = 0 s/mm² for distortion correction.

All T1-weighted images were visually inspected for image quality and motion artifacts prior to preprocessing. Structural images were processed using established neuroimaging pipelines implemented in FSL (FMRIB Software Library). T1-weighted images were cropped at the C1 vertebral level to exclude the neck region and reduce segmentation errors. Following brain extraction, images were spatially normalized to the MNI152 template and segmented into gray matter (GM), white matter (WM), and cerebrospinal fluid (CSF). Gray matter images were nonlinearly registered to a study-specific template, and modulation was applied to preserve local gray matter volume following spatial normalization, consistent with standard voxel-based morphometry procedures.

Diffusion-weighted images underwent preprocessing, including B0-field distortion correction, eddy-current correction, skull stripping, tensor fitting, and calculation of fractional anisotropy (FA) maps. FA images were corrected for brain-edge artifacts and aligned to standard MNI space using nonlinear registration. The aligned FA images were merged into a single 4D image, and a skeletonized representation of white matter tracts was generated by projecting FA values onto a mask of regions with the highest FA values.

Regional morphometric measures were extracted from cortical and subcortical regions of interest (ROIs) defined according to a standardized anatomical atlas. Total intracranial volume (TIV) was estimated for each participant and included as a covariate in subsequent structural analyses when appropriate.

### 2.4 Multilayer Network Construction

To capture the complex organization of brain structure and identify groups of patients with similar neuroanatomical profiles, multilayer brain networks were constructed following established network neuroscience frameworks [13,17–19]. In this approach, each layer represents a distinct modality or feature set derived from the structural MRI data, while nodes correspond to individual patients characterized by their structural MRI profiles (Figure 1).

Conceptually, patients are grouped according to the similarity of their combined gray and white matter structural MRI profiles. Patients with similar neuroanatomical patterns are therefore expected to cluster together, allowing the identification of data-driven subgroups that may reflect distinct neurobiological phenotypes.

Using the MultinetX library [12] nodes were characterized by disease-related gray matter density and fractional anisotropy patterns derived after removing demographic and normative sources of variability. Demographic variables and natural occurring brain variation were regressed in 2 steps. Using scans of control subjects, an initial linear regression was fitted to each regional volumetric measure, with age and sex as covariates to calculate residuals for all subjects. Normal brain variability was extracted from a principal component analysis on control data. Allowing us to isolate the disease related changes in patients through regression and standardization.

Inter-patient associations were estimated by computing pairwise statistical relationships between atrophy maps across participants, resulting in weighted adjacency matrices for each layer, consistent with prior applications of network-based morphometric analyses.

Separate network layers were constructed to represent the different structural modalities, gray matter and white matter. Within-layer edges captured similarities between atrophy maps within the same tissue type, whereas inter-layer connections linked corresponding nodes across layers, thereby preserving node identity across modality. Inter-layer coupling was defined using identity links between patients, following standard multilayer network formalism [13], to ensure a single individual would not be assigned to different clusters due to differences in tissue type.

The resulting multilayer network architecture allows the integration of regional structural information across layers while maintaining their distinct contributions. This approach has been shown to be particularly suitable for modeling brain organization across multiple scales or conditions and for capturing interactions that are not accessible using single-layer networks [11,19].

### 2.5 Clustering Analysis

A Louvain community detection algorithm was first applied to the multilayer network across multiple iterations to identify stable patterns of patient similarity. The resulting community assignments were combined into a consensus matrix reflecting the proportion of times that pairs of patients were assigned to the same community. Agglomerative hierarchical clustering using Ward’s linkage criterion was subsequently applied to the normalized consensus matrix. The Silhouette Coefficient was used to evaluate cluster separability and support selection of the final clustering solution.

The resulting clusters were subsequently characterized using voxel-based morphometry, cognitive performance measures, and health-related quality of life outcomes. Post hoc analyses were conducted to assess differences between clusters and healthy controls, as well as between clusters themselves. When applicable, multiple comparisons were corrected using the false discovery rate (FDR) procedure (Figure 1).

### 2.6 Longitudinal Analysis

To examine changes over time, cognitive and clinical data were obtained at two timepoints: baseline (TP1) and six-month follow-up (TP2). Only patients who completed both assessments were included in longitudinal analyses.

For each cognitive and clinical variable, scores were adjusted for age, sex, and years of education using linear regression models, and residualized values were used for all analyses.

Longitudinal comparisons were conducted by evaluating differences between TP1 and TP2 in the entire patient group, as well as separately within each patient subgroup (Cluster 1 and Cluster 2), in order to characterize group-specific patterns across time.

Variables included cognitive performance measures derived from the CCAS-S and the Creyos Cognitive Battery, as well as health-related quality of life (SF-36), alongside morphometric brain measures obtained from structural MRI.

### 2.7 Brain–Behavior Correlations

To investigate the relationship between structural brain changes and cognitive changes over time, brain–behavior correlation analyses were performed using longitudinal change scores (Δ = TP2 – TP1) for both imaging and cognitive variables.

Analyses focused on regions of interest (ROIs) that showed significant group or cluster differences in the structural analyses. Associations between changes in brain morphology and changes in cognitive performance were assessed using partial correlation analyses, controlling for age, sex, and years of education. Correlations were computed using Pearson’s or Spearman’s coefficients depending on data distribution.

Cognitive variables included measures derived from the CCAS-S, the Creyos Cognitive Battery, and the SF-36 questionnaire. Multiple comparisons were controlled using false discovery rate (FDR) correction where appropriate. These analyses aimed to determine whether longitudinal structural brain changes were associated with concurrent changes in cognitive and clinical functioning in patients with Long COVID.

### 2.8 Statistical Analysis

Statistical analyses were performed using R software version 4.4.1. Data normality was assessed using the Shapiro–Wilk test. Prior to cross-sectional cognitive analyses, cognitive variables were adjusted for age, sex, and years of education using linear regression models. Residual values from these models were extracted and used in subsequent analyses to minimize the influence of demographic factors on cognitive performance.

Cross-sectional group comparisons were performed using independent-samples t-tests or Mann– Whitney U tests depending on data distribution. Planned pairwise comparisons were conducted between healthy controls and Cluster 1, healthy controls and Cluster 2, and between Cluster 1 and Cluster 2.

Categorical variables were analyzed using Fisher’s exact tests when expected cell counts were small. Odds ratios (ORs) were calculated for comparisons of clinical symptom frequencies between patient subgroups.

Structural neuroimaging group comparisons were performed using general linear models implemented in FSL, including age and total intracranial volume (TIV) as covariates where appropriate.

Pearson’s or Spearman’s correlation coefficients were used according to data distribution. When applicable, multiple comparisons were corrected using the false discovery rate (FDR) procedure. Statistical significance was set at p < 0.05. Results surviving FDR correction are reported as significant, whereas results with uncorrected p < 0.05 that did not survive FDR correction are reported as trend-level findings.

## 3. RESULTS

### 3.1 Participants

Demographic and clinical characteristics are summarized in Table 1 and 2. No significant differences were observed between Long COVID patients and healthy controls in age, sex distribution, or years of education. At TP2, 32 patients completed the MRI and clinical follow-up assessments, while 31 completed the Creyos battery.

**Table 1.**
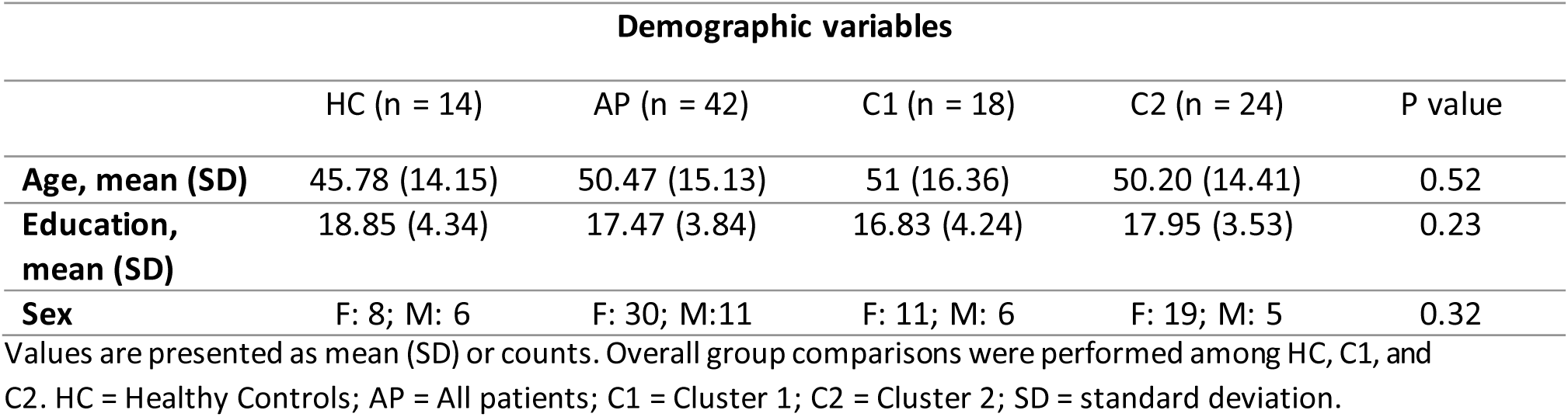
Demographic variables.

| Demographic variables |  |  |  |  |  |
| --- | --- | --- | --- | --- | --- |
|  | HC (n = 14) | AP (n = 42) | C1 (n = 18) | C2 (n = 24) | P value |
| **Age, mean (SD)** | 45.78 (14.15) | 50.47 (15.13) | 51 (16.36) | 50.20 (14.41) | 0.52 |
| **Education, mean (SD)** | 18.85 (4.34) | 17.47 (3.84) | 16.83 (4.24) | 17.95 (3.53) | 0.23 |
| **Sex** | F: 8; M: 6 | F: 30; M:11 | F: 11; M: 6 | F: 19; M: 5 | 0.32 |

Values are presented as mean (SD) or counts. Overall group comparisons were performed among HC, C1, and C2. HC = Healthy Controls; AP = All patients; C1 = Cluster 1; C2 = Cluster 2; SD = standard deviation.

**Table 2.** Clinical characteristics of study participants.

|  | AP (n = 41) | C1 (n = 17) | C2 (n = 24) | OR | P value |
| --- | --- | --- | --- | --- | --- |
| **Day since infection, mean (SD)** | 404.5 (281.4) | 411.2 (303.3) | 399.8 (271.3) |  | 0.57 |
| **1 infection, n (%)** | 31 (75.7) | 12 (70.5) | 19 (79.1) |  |  |
| **2 infection, n (%)** | 6 (14.6) | 3 (17.7) | 3 (12.5) |  |  |
| **3 infection, n (%)** | 3 (7.3) | 2 (11.8) | 1 (4.2) |  |  |
| **4 infection, n (%)** | 1 (2.4) | 0 (0) | 1 (4.2) |  |  |
| **Severity** |  |  |  |  | 0.36 |
| **0, n (%)** | 1 (2.4) | 0 (0) | 1 (4.2) |  |  |
| **1, n (%)** | 11 (26.8) | 6 (35.2) | 5 (20.8) |  |  |
| **2, n (%)** | 24 (58.5) | 10 (58.9) | 14 (58.3) |  |  |
| **3, n (%)** | 2 (4.8) | 0 (0) | 2 (8.3) |  |  |
| **4, n (%)** | 2 (4.8) | 1 (5.9) | 1 (4.2) |  |  |
| **5, n (%)** | 1 (2.4) | 0 (0) | 1 (4.2) |  |  |
| **Mood problems, n (%)** | 16 (39) | 7 (41.1) | 9 (37.5) | 0.86 | 1 |
| **Foggy mind, n (%)** | 36 (87.8) | 16 (94.1) | 20 (83.3) | 0.32 | 0.38 |
| **Confusion, n (%)** | 21 (51.2) | 10 (58.9) | 11 (45.8) | 0.6 | 0.53 |
| **Insomnia, n (%)** | 17 (41.4) | 7 (41.1) | 10 (41.6) | 1.02 | 1 |
| **Dizziness, n (%)** | 20 (48.7) | 9 (52.9) | 11 (45.8) | 0.76 | 0.75 |
| **Headaches, n (%)** | 24 (58.5) | 9 (52.9) | 15 (62.5) | 1.47 | 0.74 |
| **Fatigue, n (%)** | 35 (85.3) | 13 (76.4) | 22 (91.6) | 3.28 | 0.21 |
| **Difficulty remembering information, n (%)** | 36 (87.8) | 17 (100) | 19 (79.1) | 0 | 0.06 |
| **Anosmia, n (%)** | 10 (24.3) | 5 (29.4) | 5 (20.8) | 0.64 | 0.71 |
| **Find it easy to forget information, n (%)** | 33 (80.4) | 14 (82.3) | 19 (79.1) | 0.82 | 1 |

Values are presented as mean (SD) or *n* (%), as appropriate. P values correspond to comparisons between Cluster 1 and Cluster 2. Odds ratios (OR) represent the association between cluster membership and the presence of each clinical characteristic. Clinical information was available for 41 of the 42 patients. AP = All patients; C1 = Cluster 1; C2 = Cluster 2.

### 3.2 Clustering Analysis

Given features derived from the multilayer network, hierarchical clustering produced a dendrogram of patients. Evaluation using the silhouette score indicated that a two-cluster solution (Cluster 1 and Cluster 2) provided the highest data separability, reflecting distinct neuroanatomical profiles (Figure 2). These clusters were subsequently used for comparisons of structural brain measures, cognitive performance, and healthy controls.

**Figure 2.**
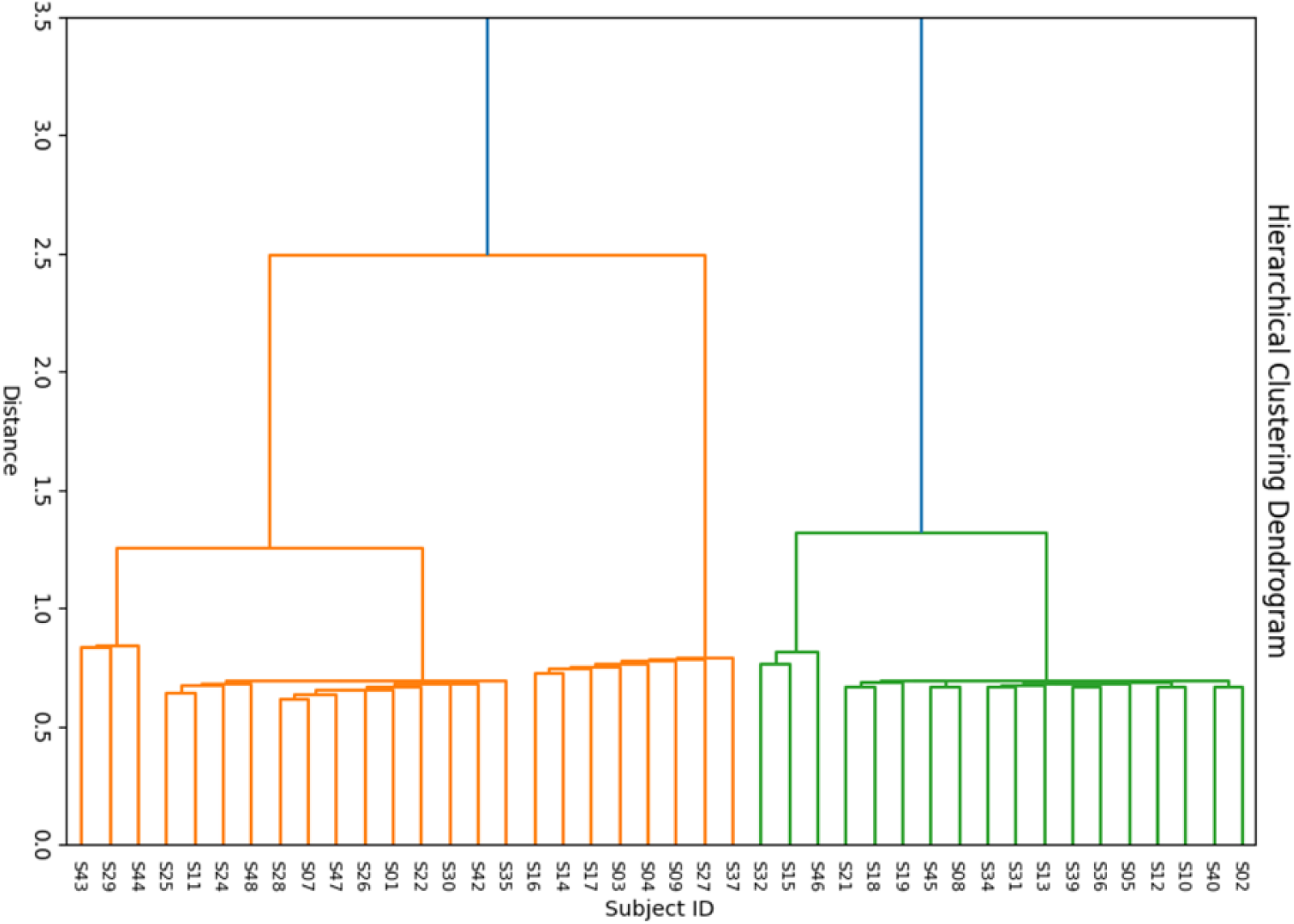
Multilayer network–based clustering of patients with Long COVID. Dendrogram illustrating the hierarchical clustering solution derived from the consensus matrix of multilayer network features. Each leaf represents an individual participant, and branch lengths reflect dissimilarity between subjects based on their structural brain profiles. Two main clusters were identified (Cluster 1 and Cluster 2), corresponding to distinct neuroanatomical patterns. Cluster assignment was determined using agglomerative hierarchical clustering applied to the consensus matrix obtained from iterative Louvain community detection. The optimal number of clusters was supported by cluster quality indices, including thesilhouette coefficient. Colors indicatecluster membership (Cluster 1: green; Cluster 2: orange).

### 3.3 Cross-sectional analyses

#### 3.3.1 Voxel-based morphometry comparison

Compared with healthy controls, Cluster 2 showed reduced gray matter density in the putamen and cerebellar lobules VIIIa and VIIIb (Figure 3a). In contrast, no significant differences were observed between healthy controls and Cluster 1. Direct comparisons between Cluster 1 and Cluster 2 revealed widespread differences predominantly involving cerebellar regions (Figure 3b).

**Figure 3.**
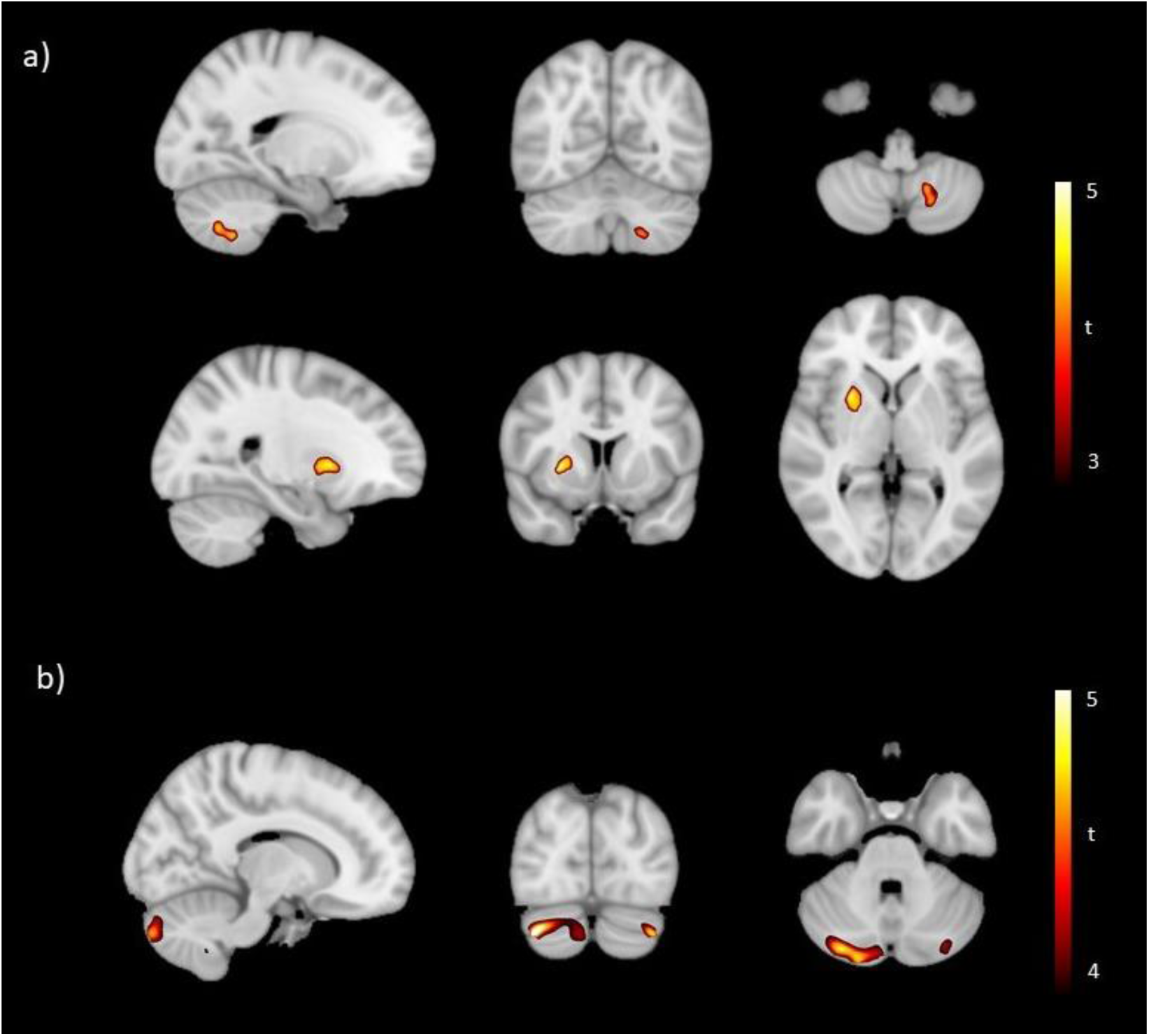
Voxel-based morphometry (VBM) group comparisons. (a) Comparison between Cluster 2 and healthy controls showing reduced gray matter density in the putamen and cerebellar lobules VIIIa and VIIIb. (b) Direct comparison between Cluster 1 and Cluster 2 revealing cluster-specific differences predominantly involving cerebellar regions. Statistical maps are displayed at the voxel level and thresholded using TFCE correction (p < 0.05, corrected). Color bars represent t-statistics. Images are shown in MNI space.

#### 3.3.2 Cognitive performance

##### CCAS-S

Compared with healthy controls, Cluster 1 showed lower performance on the Go/No-Go task (p = 0.01, d = 0.93). Cluster 2 differed from healthy controls on semantic fluency (p = 0.004, d = 1.01) and Go/No-Go performance (p = 0.037, r = 0.33). No significant differences were observed between Cluster 1 and Cluster 2. None of these comparisons survived FDR correction (Table 3).

**Table 3.** Cross-sectional differences in cognitive performance and health-related quality of life between healthy controls and Long COVID subgroups.

| Variable | Groups | Test statistic | p | $p_{FDR}$ | Effect size |
| --- | --- | --- | --- | --- | --- |
| **SCCA** |  |  |  |  |  |
| Go/No-Go | HC vs C1 | 2.61 | 0.01 | ns | 0.93 |
| Semantic fluency | HC vs C2 | 3.02 | 0.004 | ns | 1.01 |
| Go/No-Go | HC vs C2 | 237 | 0.03 | ns | 0.33 |
| **Creyos battery** |  |  |  |  |  |
| Spatial planning score | HC vs C1 | 186 | 0.02 | ns | 0.40 |
| Spatial Planning correct answers | HC vs C1 | 2.60 | 0.01 | ns | 0.92 |
| Odd One Out correct answers | HC vs C1 | 2.04 | 0.04 | ns | 0.73 |
| Feature match correct answers | HC vs C2 | 2.31 | 0.02 | ns | 0.77 |
| Polygons correct answers | HC vs C2 | 2.50 | 0.01 | ns | 0.84 |
| Spatial planning correct answers | HC vs C2 | 2.14 | 0.03 | ns | 0.72 |
| Token search duration time span | HC vs C2 | -2.82 | 0.007 | ns | -0.95 |
| Spatial span correct answers | C1 vs C2 | 321 | 0.006 | ns | 0.41 |
| Polygon errors made | C1 vs C2 | 2.67 | 0.01 | ns | 0.83 |
| **SF-36** |  |  |  |  |  |
| General health | HC vs C1 | 3.47 | 0.001 | 0.003 | 1.23 |
| Emotional well being | HC vs C1 | 2.18 | 0.03 | 0.04 | 0.77 |
| Energy fatigue | HC vs C1 | 5.58 | <0.001 | <0.001 | 1.98 |
| Limitations emotional problems | HC vs C1 | 3.49 | 0.001 | 0.003 | 1.24 |
| Limitations physical health | HC vs C1 | 5.13 | <0.001 | <0.001 | 1.83 |
| Physical functioning | HC vs C1 | 2.22 | 0.03 | 0.04 | 0.79 |
| General health | HC vs C2 | 4.08 | <0.001 | <0.001 | 1.37 |
| Limitations physical health | HC vs C2 | 5.17 | <0.001 | <0.001 | 1.74 |
| Pain | HC vs C2 | 2.50 | 0.01 | 0.02 | 0.84 |
| Physical functioning | HC vs C2 | 4.42 | <0.001 | <0.001 | 1.41 |
| Energy fatigue | HC vs C2 | 279.5 | <0.001 | 0.001 | 0.54 |

Values correspond to statistically significant pairwise comparisons identified in the cross-sectional analyses. Test statistics are reported according to the statistical test applied. p values were adjusted for multiple comparisons using the Benjamini–Hochberg false discovery rate (FDR) procedure. HC = healthy controls; C1 = Cluster 1; C2 = Cluster 2; ns = not significant after FDR correction.

##### Creyos Battery

Compared with healthy controls, Cluster 1 showed lower performance on Odd One Out correct answers (p = 0.04, d = 0.70) and Spatial Span performance, including correct answers (p = 0.01, d = 0.92) and total score (p = 0.02, r = 0.4).

Cluster 2 showed reduced performance compared with healthy controls on Feature Match correct answers (p = 0.02, d = 0.77), Polygon correct answers (p = 0.01, d = 0.84), Spatial Span correct answers (p = 0.03, d = 0.72), and Token task time (p = 0.007, d = −0.95).

Direct comparisons between clusters revealed differences in Polygon errors (p = 0.01, d = 0.83) and Spatial Span correct answers (p = 0.006, r = 0.41).

Although none of these comparisons survived FDR correction, several measures demonstrated medium-to-large effect sizes (d = 0.70–1.01), indicating that the magnitude of between-group differences was not negligible despite the lack of corrected statistical significance (table 3).

##### Functional status (SF-36)

Compared with healthy controls, Cluster 1 showed significantly lower scores across six SF-36 domains, including Energy/Fatigue (p < 0.001, d = 1.99), Role limitations due to physical health (p < 0.001, d = 1.83), General Health (p = 0.003, d = 1.24), Role limitations due to emotional problems (p = 0.003, d = 1.25), Emotional well-being (p = 0.04, d = 0.78), and Physical Functioning (p = 0.04, d = 0.79).

Cluster 2 also showed significantly lower scores compared with healthy controls in five domains, including General Health (p < 0.001, d = 1.37), Physical Functioning (p < 0.001, d = 1.41), Role limitations due to physical health (p < 0.001, d = 1.74), Energy/Fatigue (p = 0.001, d = 0.55), and Pain (p = 0.02, d = 0.84).

No significant differences were observed between Cluster 1 and Cluster 2 (table 3).

### 3.4 Longitudinal analyses

#### 3.4.1 Voxel-based morphometry comparison

Longitudinal VBM analyses in the patient group revealed a significant increase in gray matter density from TP1 to TP2 in the medial frontal cortex. No other regions showed significant longitudinal changes. No significant longitudinal effects were observed between the two patient clusters (Figure 3).

**Figure 4.**
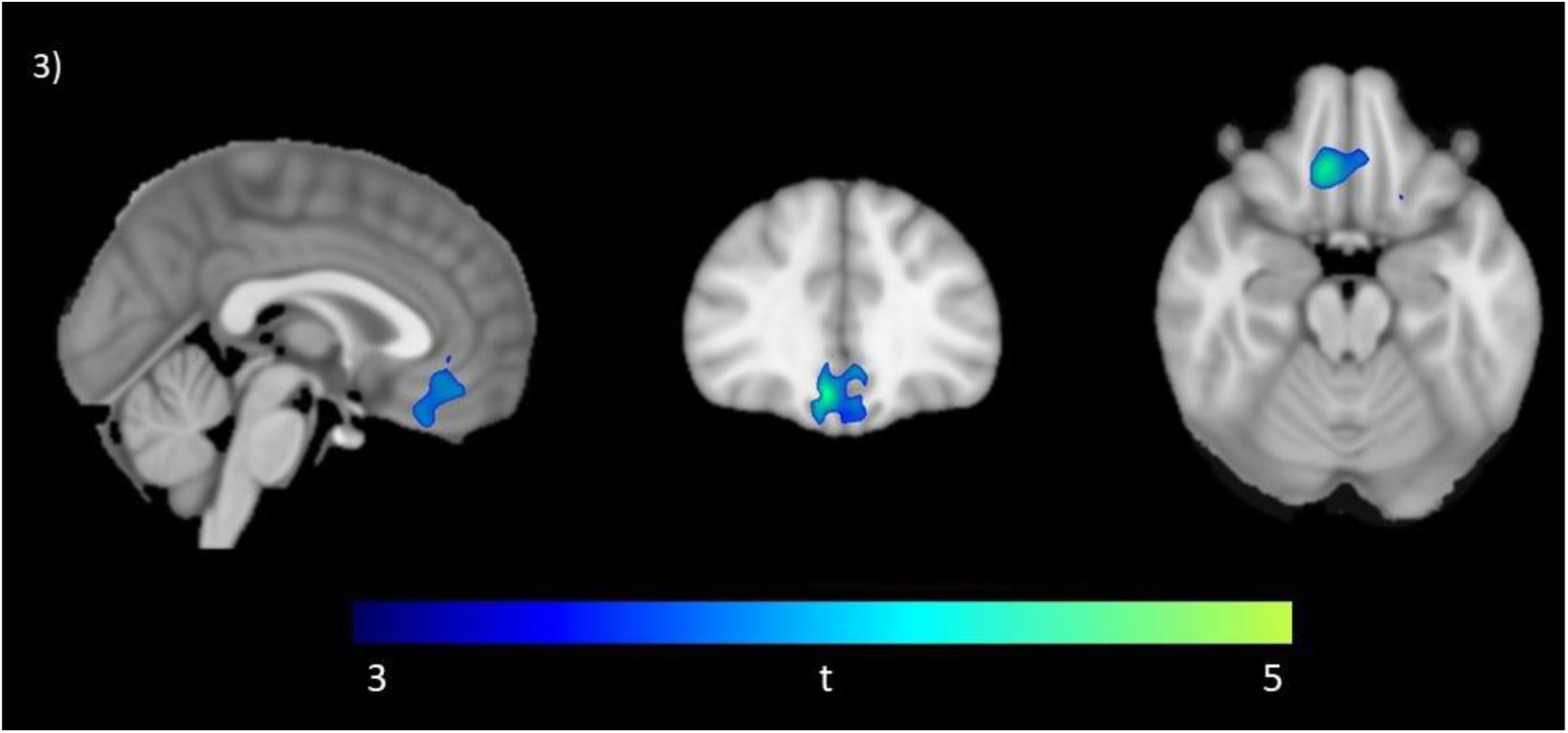
Longitudinal voxel-based morphometry (VBM) results. Group-level comparison between Timepoint 1 and 2 showing a significant increase in gray matter density in the medial frontal cortex in patients with Long COVID. No additional regions showed significant longitudinal changes. Statistical maps are displayed at the voxel level and thresholded using TFCE correction (p < 0.05, corrected). Color bar represents t-statistics. Images are shown in MNI space.

#### 3.4.2 Cognitive performance

##### CCAS-S

Cluster 2 showed a trend-level longitudinal change in the Cube task (p = 0.04, d = −0.49). No significant longitudinal changes were observed at the global level or within Cluster 1 (table 4).

**Table 4.** Longitudinal changes in cognitive performance between TP1 and TP2 in Long COVID patients.

| Variable | Group | Test statistic | p | $p_{FDR}$ | Effect size |
| --- | --- | --- | --- | --- | --- |
| **SCCA** |  |  |  |  |  |
| Cube | C2 | -2.12 | 0.04 | ns | -0.49 |
| **Creyos battery** |  |  |  |  |  |
| Spatial planning errors made | Patient group | -2.79 | 0.007 | ns | -0.48 |
| Spatial span correct answers | C1 | 2.17 | 0.04 | ns | 0.73 |
| Spatial span duration time span | C1 | 2.11 | 0.04 | ns | 0.60 |
| Spatial planning errors made | C2 | -2.06 | 0.04 | ns | -0.51 |

Values correspond to statistically significant within-group longitudinal comparisons between TP1 and TP2. Test statistics are reported according to the statistical test applied (paired Student's *t* test or Wilcoxon signed-rank test). *p* values were adjusted using the Benjamini–Hochberg false discovery rate (FDR) procedure. Patient group = all patients; C1 = Cluster 1; C2 = Cluster 2; ns = not significant after FDR correction.

##### Creyos Battery

At the level of the total patient group, a trend-level change was observed for Spatial Span errors made (p = 0.007, d = −0.48). Within Cluster 1, longitudinal trends were observed for Spatial Span correct answers (p = 0.04, d = 0.73) and Spatial Span time (p = 0.04, d = 0.60). Cluster 2 showed a trend-level change in Spatial Span errors made (p = 0.045, d = −0.51) (table 4).

##### Functional status (SF-36)

No significant longitudinal changes were observed in SF-36 domains at the level of the total patient group or within either cluster.

### 3.5 Brain–behavior correlations

Correlations between longitudinal changes in medial frontal cortex gray matter density and changes in cognitive and functional measures were examined using delta values (TP2 − TP1).

No significant correlations were observed between medial frontal cortex changes and CCAS-S delta scores at the level of the entire patient group or within clusters.

For the Creyos battery, several nominally significant correlations were observed between longitudinal changes in medial frontal gray matter density and cognitive performance; however, none survived correction for multiple comparisons. At the whole-group level, changes in medial frontal gray matter density were associated with Polygon score (p = 0.01, r = 0.45) and Polygon correct answers (p = 0.03, r = 0.39). Within Cluster 1, changes in medial frontal gray matter density were associated with Double Trouble errors made (p = 0.006, r = 0.78) and Polygon score (p = 0.03, r = 0.66). No significant correlations were observed in Cluster 2.

Regarding functional outcomes, medial frontal cortex changes showed a trend correlation with changes in the Emotional Well-being domain of the SF-36 at the level of the entire patient group (p = 0.03, r = 0.38), with no significant correlations observed within individual clusters.

## 4. DISCUSSION

The present study provides converging evidence that post-COVID cognitive symptoms are not associated with a uniform neuroanatomical profile. Using a data-driven clustering approach, we identified two distinct patient subgroups characterized by differential patterns of brain structure, supporting the existence of separable neuroanatomical phenotypes in Long COVID. This finding is consistent with previous work highlighting substantial interindividual variability in symptom presentation and brain alterations following SARS-CoV-2 infection [9,10]. Notably, group differences were more pronounced in measures of brain structure and health-related quality of life, whereas cognitive differences were more subtle and generally did not survive correction for multiple comparisons. Nevertheless, several cognitive measures showed medium-to-large effect sizes despite not surviving correction for multiple comparisons. This pattern suggests that potentially meaningful cognitive differences may exist between neuroanatomically defined subgroups but could remain undetected after stringent multiple-comparison correction in a modestly sized sample [20]. Overall, these results support the view that Long COVID comprises multiple neurobiological phenotypes rather than a single homogeneous entity [3,8], underscoring the relevance of subtyping approaches for advancing mechanistic understanding and informing more personalized clinical strategies.

### 4.1 Structural Brain Findings

Structural differences between patient clusters were primarily characterized by reduced gray matter density in cerebellar lobules VIIIa and VIIIb and in the putamen. These findings align with accumulating evidence implicating both cerebellar and subcortical structures in the cognitive manifestations of Long COVID [5,7,8]. Beyond its established role in motor coordination, the cerebellum—particularly posterior lobules such as VIIIa and VIIIb—has been increasingly recognized as a key contributor to executive control, working memory, and visuospatial processing [21–23]. Alterations in these regions may therefore provide a plausible neural substrate for the executive and visuospatial difficulties reported in a subset of patients with persistent post-COVID cognitive symptoms.

Similarly, involvement of the putamen is consistent with its role in fronto-striatal circuits supporting cognitive flexibility, attentional control, and action selection [24]. Structural and functional abnormalities of the putamen have been described in neurological and neuropsychiatric conditions characterized by cognitive slowing and executive dysfunction, and emerging evidence suggests that basal ganglia structures may also be vulnerable in Long COVID [3,8]. Together, these findings support the notion that cognitive symptoms in Long COVID may arise from disruptions in distributed subcortical–cortical networks rather than isolated regional damage. Such distributed patterns are more readily captured by approaches that consider large-scale brain organization.

Longitudinal analyses revealed a significant increase in gray matter density in the medial frontal cortex over the six-month follow-up period at the group level. This region plays a central role in executive control, attentional regulation, performance monitoring, and cognitive effort, and has been consistently implicated in conditions characterized by cognitive fatigue and reduced processing efficiency [25,26]. Similar medial frontal alterations have also been reported in previous neuroimaging studies of Long COVID, supporting its involvement in the context of persistent post-infectious cognitive symptoms [3,5].

This increase in gray matter density may reflect compensatory or plastic reorganization processes during recovery; however, alternative interpretations should be considered [27]. Voxel-based morphometry is sensitive to multiple biological mechanisms, including synaptic density changes, neuroinflammation, and fluid shifts, and therefore changes in gray matter density should not be interpreted as direct evidence of neuronal recovery [28]. Consequently, the observed longitudinal changes may reflect a range of biological processes whose underlying mechanisms cannot be determined from VBM alone. Importantly, this structural change was not accompanied by robust improvements in objective cognitive performance at the group level, suggesting a dissociation between structural and functional trajectories over time. The absence of significant longitudinal effects within clusters further highlights heterogeneity in recovery patterns and may also reflect limited statistical power for subgroup analyses.

### 4.2 Cognitive Findings

Despite persistent subjective cognitive complaints, longitudinal analyses did not reveal robust improvements or declines in objective cognitive performance over the six-month follow-up. At the group level, performance on both the CCAS-S and Creyos battery remained largely stable, with only trend-level changes observed in selected visuospatial and executive measures. This pattern suggests relative stability of objective cognitive functioning over time, despite ongoing symptom perception.

This stability may reflect multiple, non-mutually exclusive mechanisms. First, cognitive impairment in Long COVID may be subtle or subclinical and therefore not fully captured by standard neuropsychological instruments over relatively short follow-up intervals [16,29]. Second, prior studies have described a dissociation between subjective cognitive complaints and objective performance in Long COVID, particularly in attention, processing speed, and executive domains [30,31], suggesting that “brain fog” may be more closely related to cognitive effort, fatigue, or affective symptoms than to measurable deficits.

The trend-level longitudinal effects observed in visuospatial working memory tasks, such as Spatial Span, are nonetheless noteworthy. These domains have been repeatedly implicated in post-COVID cognitive dysfunction and are known to be sensitive to subtle alterations in fronto-cerebellar and fronto-striatal networks [7,32]. Although these effects did not survive correction for multiple comparisons, their consistency across analyses suggests that visuospatial and executive domains may represent particularly vulnerable cognitive targets in Long COVID and warrant further investigation in larger longitudinal cohorts.

Although between-cluster cognitive differences were modest and did not survive correction for multiple comparisons, several executive and visuospatial measures showed medium-to-large effect sizes, suggesting that neuroanatomically defined subtypes may exhibit distinct cognitive profiles. The observed medium-to-large effect sizes suggest that the magnitude of the differences was not negligible and could reflect limited statistical power associated with the relatively small subgroup sizes rather than the absence of meaningful cognitive differences. Replication in larger cohorts will be necessary to determine the robustness and clinical relevance of these findings. Importantly, structural stratification remained evident even in the absence of robust behavioral differentiation, supporting the notion that neurobiological heterogeneity may not always be fully reflected in conventional cognitive measures.

The absence of significant differences between Cluster 1 and healthy controls does not necessarily indicate the absence of clinically relevant dysfunction in this subgroup. Rather, it may reflect subtle or compensable neural alterations not captured by conventional morphometric or cognitive measures, despite persistent subjective symptoms such as “brain fog.” This dissociation between subjective complaints and objective performance has been consistently reported in post-COVID populations and may reflect altered cognitive effort, fatigue-related mechanisms, or network-level inefficiencies rather than focal structural damage [16,30,31].

### 4.3 Brain–Behavior Relationships

Beyond group-level effects, exploratory brain–behavior analyses provided additional insight into the potential functional relevance of these structural changes. Longitudinal increases in medial frontal gray matter density were nominally associated with performance on selected visuospatial and executive measures, although these associations did not survive correction for multiple comparisons. The medial frontal cortex is a key hub for cognitive control, working memory, response inhibition, and performance monitoring [25,33], functions that are frequently affected in Long COVID.

Nominal associations with the Interlocking Polygons task suggest that medial frontal structural changes may relate to visuospatial integration and planning processes. Altho ugh exploratory and not significant after correction for multiple comparisons, these findings are consistent with the hypothesis that longitudinal structural reorganization may have functional relevance during recovery. Similarly, the nominal association with the Double Trouble task observed in Cluster 1 is consistent with the established role of medial frontal regions in conflict monitoring, inhibitory control, and executive regulation across cognitive domains [33,34]. Together, these exploratory findings highlight the potential cognitive relevance of medial frontal alterations while underscoring the complexity of linking regional structural changes to specific cognitive processes.

### 4.4 Interpreting Heterogeneity in Long COVID

Overall, the present findings suggest that post-COVID cognitive symptoms may reflect disruption of large-scale control networks rather than isolated regional deficits. Structural involvement of fronto - parietal and cerebellar regions—key components of executive control, attentional regulation, and cognitive flexibility—is consistent with models conceptualizing cognitive control as an emergent property of distributed brain systems [26,35]. The role of fronto-parietal and cingulo-opercular networks in sustaining goal-directed behavior [36], together with the modulatory contribution of the cerebellum to executive function [32,37], further supports this interpretation.

Within this framework, the observed interindividual variability reinforces the notion that post-COVID cognitive impairment is not a unitary syndrome but rather a heterogeneous condition with partially overlapping neural substrates. This variability supports the use of data-driven clustering approaches to identify neurobiologically meaningful subtypes with potentially distinct cognitive and structural profiles.

Notably, the identified subgroups were not explained by conventional demographic or clinical variables, including age, education, COVID-19 severity, time since infection, or the prevalence of persistent symptoms. This finding suggests that neuroanatomical heterogeneity in Long COVID may not be fully captured by the clinical characterization and highlights the potential value of neuroimaging-based stratification approaches for identifying biologically meaningful variability that is not captured by conventional clinical measures [9,38].

An alternative interpretation is that the identified clusters may represent different stages of recovery rather than biologically distinct neuroanatomical phenotypes. Although the present study cannot distinguish conclusively between these possibilities, several observations favor the latter interpretation. Specifically, the identified subgroups were not explained by conventional demographic or clinical variables, including time since infection, COVID-19 severity, or the prevalence of persistent symptoms. At the same time, because clustering was performed only at baseline, we cannot determine whether patients would maintain the same subgroup membership over time. Future longitudinal studies incorporating repeated clustering analyses will therefore be important to establish whether these subgroups represent stable neurobiological phenotypes or dynamic stages along the recovery process.

Consistent with this perspective, the absence of strong cognitive differences between clusters does not diminish the relevance of structural stratification. Instead, it suggests that brain alterations may precede, accompany, or be differentially compensated across individuals. From this viewpoint, approaches that explicitly model heterogeneity—rather than relying on group averages—are particularly well suited for conditions such as Long COVID, where clinical trajectories are variable and potentially non-linear.

Previous work has shown that normative modeling and subtyping approaches can reveal meaningful neurobiological structure even in the absence of clear behavioral separation [38]. Collectively, these findings support the use of neurobiological stratification strategies to refine the characterization of Long COVID and may ultimately inform more personalized clinical approaches and monitoring strategies.

## 5. LIMITATIONS

Several limitations should be considered when interpreting the present findings. First, while the present sample enabled the identification of meaningful structural differences between subgroups, larger longitudinal cohorts would provide greater statistical power to determine whether the medium-to-large cognitive effect sizes observed in the present study represent robust between-group differences or instead reflect limited statistical power associated with the modest sample size. Larger samples would also improve the ability to detect more subtle cognitive or structural changes over time and to better characterize the interindividual variability that is a defining feature of Long COVID. Second, although voxel-based morphometry (VBM) and diffusion-based measures provide complementary information about gray and white matter structure, they remain indirect markers of underlying neurobiological processes. The absence of additional imaging modalities (e.g., functional or molecular imaging) limits mechanistic interpretation of the observed findings, particularly regarding the relative contributions of neuroinflammation, vascular alterations, and synaptic plasticity. Third, the modest longitudinal associations between structural changes and cognitive performance limit stronger inferences regarding brain–behavior relationships. This dissociation may reflect compensatory mechanisms, subclinical cognitive involvement, or limited sensitivity of the cognitive measures to detect subtle changes. Finally, the data-driven clustering approach, while useful for capturing heterogeneity, is dependent on feature selection and methodological choices. Replication in larger, independent samples using longitudinal and multimodal data will be necessary to confirm and refine the identified subgroups.

## 6. CONCLUSION

In summary, these exploratory findings suggest that post-COVID cognitive symptoms are associated with heterogeneous neurobiological profiles rather than a uniform pattern of brain or cognitive impairment. Using a multilayer network approach combined with structural MRI and cognitive assessment, we identified data-driven patient subgroups with distinct neuroanatomical characteristics, supporting the view that Long COVID is unlikely to represent a single neurobiological entity. Although cognitive differences between subgroups were modest, the observed structural heterogeneity suggests that biologically meaningful variability may exist even in the absence of robust behavioral separation. These findings highlight the value of data-driven stratification for improving the characterization of Long COVID and provide a foundation for future hypothesis-driven studies aimed at refining neurobiological subtypes and informing more personalized clinical approaches.

## 9. AUTHOR CONTRIBUTIONS

- Conceptualization: A.O.R.M, B.M., J.F.R., C.R.H.C.
- Data curation: A.O.R.M, B.M., C.R.H.C
- Formal analysis: A.O.R.M, B.M.
- Funding acquisition: C.R.H.C
- Investigation: A.O.R.M, B.M., C.R.H.C.
- Methodology: A.O.R.M, B.M., C.R.H.C.
- Project administration: C.R.H.C.
- Resources: C.R.H.C
- Software: A.M.O., C.R.H.C.
- Supervision: J.F.R., C.R.H.C.
- Validation: A.O.R.M, B.M., J.F.R., A.M.O., C.R.H.C.
- Visualization: A.O.R.M, J.F.R., C.R.H.C.
- Writing – original draft: A.O.R.M.
- Writing – review and editing: A.O.R.M, B.M., J.F.R., A.M.O., C.R.H.C.

## 10. ETHICS STATEMENT

The study was approved by the institutional research ethics board of the Nova Scotia Health Authority and Dalhousie University. All procedures were conducted in accordance with the Declaration of Helsinki. Written informed consent was obtained from all participants prior to study participation.

## 11. DATA AVAILABILITY STATEMENT

Data is available for researchers who meet the criteria for access to confidential data. Access requests must be submitted in writing to the corresponding author and approval from the relevant ethics committee.

## 12. ACKNOWLEDGEMENTS

We thank all patients and volunteers for their kind and valuable participation.

